# QTL mapping in an interspecific sorghum population uncovers candidate regulators of salinity tolerance

**DOI:** 10.1101/2020.08.05.238972

**Authors:** Ashley N. Hostetler, Rajanikanth Govindarajulu, Jennifer S. Hawkins

**Author notes:** Author for correspondence: Ashley N. Hostetler 53 Campus Drive Department of Biology West Virginia University Morgantown, WV 26505.

## Abstract

Salt stress impedes plant growth and disrupts normal metabolic processes, resulting in decreased biomass and increased leaf senescence. Therefore, the ability of a plant to maintain biomass when exposed to salinity stress is critical for the production of salt tolerant crops. To identify the genetic basis of salt tolerance in an agronomically important grain crop, we used a recombinant inbred line (RIL) population derived from an interspecific cross between domesticated *Sorghum bicolor* (inbred Tx7000) and a wild relative, *Sorghum propinquum*. A high-density genetic map was generated from 177 F_3:5_ RILs and covered the 10 *Sorghum* chromosomes with 1991 markers. The genetic map was used to identify 19 total QTL related to plant growth and overall health in optimal and saline conditions. Of these 19 QTL detected, 10 were specific to the salt stress response. The salt-responsive QTL contained numerous genes that have been previously shown to play a role in ionic tolerance, tissue tolerance, and osmotic tolerance, including many aquaporins.

## Introduction

Soil salinity imposes abiotic stress on plants when soluble ions, such as Na^+^ and Cl^−^, accumulate in the soil surrounding the root rhizosphere. Saline soils affect more than 6% of total land and 20% of irrigated land ^1–4^, and while most of the lands with elevated salinity have arisen from natural causes, anthropogenic factors have recently led to the increase of salinity in lands cultivated for agriculture ^5^. Agricultural production is threatened by soil salinity because crops that are most affected include grain crops. Grain crops such as wheat, rice, maize, barley, millet, and sorghum are typically glycophytic ^5–7^, therefore soil salinity causes significant reductions in plant biomass and yield; however, crop tolerance varies among species and genotypes ^5^. Among monocots, rice is the most sensitive ^8^, whereas barley can tolerate almost twice as much NaCl before experiencing a comparable loss in dry matter ^9^. Furthermore, at the genotype level, maize lines show variation in tolerance to foliar sodium ^10,11^.

During salt stress, reductions in plant biomass and yield are due to both ion-independent or ion-dependent factors ^5,12–18^. When salts initially accumulate in the soil, the osmotic potential of the soil-water decreases, resulting in a reduction of water extracted by plant roots. This osmotic stress causes a sudden, short term loss of water, cell volume, and turgor from leaf cells and results in a reduction of both above and belowground growth ^5^. During the subsequent ionic phase of salinity stress, ions accumulate to toxic concentrations and disrupt normal metabolic processes and ionic homeostasis. Specifically, the accumulation of Na^+^ disrupts the uptake and distribution of K^+^, which is essential for protein synthesis ^19,20^, enzymatic reactions ^21^, and signaling ^22^. Ionic stress results in the inactivation of enzymes and the generation of reactive oxygen species ^23–25^, which results in increased leaf senescence ^5^.

Sorghum (*Sorghum bicolor* L. Moench), a staple crop for food, fuel, and feed production ^26,27^, is notable for its varieties that are naturally drought and salt tolerant ^28–37^. In a previous study, we measured the variation in salinity tolerance across a diverse panel of *Sorghum* genotypes that included wild species, domesticated *S. bicolor* landraces, and improved *S. bicolor* lines ^37^. Salinity tolerance was assessed as the maintenance of biomass following 12 weeks of 75 mM salt exposure (sodium chloride, NaCl). After long-term exposure to salinity, *S. bicolor* genotypes maintained between 30% - 95% of their biomass compared to genotypes in non-saline (control) conditions. The genotype that had the greatest reduction in biomass in response to salt exposure was the wild species *S. propinquum*, which maintained only 5% of its aboveground biomass ^37^. The observed variation in biomass retention upon exposure to saline conditions ^37^ indicates that there is quantitative genetic variation in salinity tolerance in *Sorghum*. Specifically, the findings from our previous study set the foundation for the parental genotypes of the recombinant inbred line (RIL) population used here.

In the work presented here, a RIL population constructed from a cross between *S. propinquum* (95% biomass reduction) and *S. bicolor* (Tx7000 - landrace *durra*; 5% biomass reduction) was used to dissect the genetic underpinnings of salinity tolerance. We developed a high-density genetic map from 177 F_3:5_ RILs and identified quantitative trait loci (QTL) associated with biomass-related traits during salt exposure. Additionally, these findings and this population establish a resource that can be used to further dissect the underlying genetic basis of osmotic tolerance, ionic tolerance, and tissue tolerance.

## Materials and Methods

### Plant Material

A RIL mapping population derived from an interspecific cross of *S. propinquum* and *S. bicolor* (inbred Tx7000, landrace *durra*) was used to investigate the genetic underpinnings in salinity tolerance. The RIL population consists of 177 F_3:5_ lines with 75% (132 RILs) of the individuals being F_5_, 18% (31 RILs) of the individuals being F_4_, and 7% (14 RILs) of the individuals being F_3_. Each line was derived as described in Govindarajulu et al. (2021) by the single seed descent method ^38,39^.

### Experimental Conditions

In a controlled greenhouse room, five seeds of each RIL were planted in 5 cm x 5 cm x 5 cm planting plugs filled with metromix soil. Target germination conditions were 21°C, 75% humidity, and 4.5 vapor pressure deficit (VPD). During germination, seedlings were regularly misted with non-saline tap water and watered with a 20-10-20 N-P-K fertilizer (J.R. Peters, Inc., Allentown, PA, USA) diluted to 200 mg N L^−1^ every 4^th^ day. Once all plants reached at least the third leaf stage (V3) of development (approximately 32 days post-planting), soil plugs were transplanted into 5 cm x 5 cm x 25 cm treepots (Stuewe and Sons, Tangent, OR, USA) filled with silica sand #4. Target growth conditions were 27°C day/23°C night with 16 h ambient daylight and 75% humidity. Following transplant, seedlings were watered to saturation with non-saline tap water daily for two weeks to provide a period of establishment. All plants were fertilized twice weekly with a 20-10-20 N-P-K fertilizer (J.R. Peters, Inc., Allentown, PA, USA) diluted to 200 mg N L^−1^ for the remainder of the study. Following establishment, three of the five biological replicates were randomly assigned to salt treatment and two of the five biological replicates were randomly assigned to a control treatment. All seedlings (two control plants and three treatment plants per line) were arranged in a randomized design. Given the variation in *Sorghum* salinity tolerance ^37,40–43^ and our previous findings ^37^, an extended 75 mM NaCl treatment was deemed appropriate for identifying salt tolerant and sensitive genotypes. Plants were watered once daily, in accordance with their treatment assignment (75 mM NaCl or 0 mM NaCl), to complete saturation for the duration of the experiment. Treatment began 51 days after planting and plants were treated for a total of 45 days.

### Phenotypic Measurements

The following phenotypes were measured 45 days after treatment began: height (cm), rank score, root biomass (g), dead aboveground biomass (g), live aboveground biomass (g), total aboveground biomass (g), and total biomass (g). Height (cm) was taken from the base of the stem to the tip of the newest emerged leaf. Rank score was a qualitative score that described overall leaf ‘greenness’, leaf health, and mortality, where plants that displayed no signs of stress received a low rank score, and plants that were extremely stressed or had died received a high rank score (**Table 1**). Rank score was assessed by the same person to minimize bias. All biomass measurements were taken on tissue collected from a destructive harvest and dried in 65°C for a minimum of 72 h. Root biomass (g) was the total belowground biomass collected. Roots were rinsed in water to remove all dirt and sand prior to drying. Dead aboveground biomass (g) included all biomass (leaves, tillers, and/or stem) attached to the plant that was more than 50% brown; whereas live aboveground biomass (g) included all biomass attached to the plant that was more than 50% green. Total aboveground biomass (g) was the sum of live and dead aboveground biomass, while total biomass (g) included live, dead, and root biomass. Mortality was scored as 1 if plants were alive and 0 if plants were dead.

**Table 1.** Rank scoring parameters of plant vigor. Plant vigor was assessed on a scale of 0 to 5 with 0 indicating no signs of stress and 5 indicating plant death.

| Score | Observation |
| --- | --- |
| 0 | No leaf signs |
| 1 | Some leaf and leaf tip curling |
| 2 | Severe leaf and leaf tip curling, few leaves elongated |
| 3 | Most leaves dead but still producing new leaves |
| 4 | Plant still alive but no new growth |
| 5 | Plant dead |

The stress tolerance index (STI) is a valuable metric when comparing genotypic tolerance within a population ^15^. The STI value for each trait was calculated using the following formula, where Y is the phenotypic trait, control is the value of the trait in 0 mM NaCl conditions, salt is the value of the trait in 75 mM NaCl conditions, and control average is the population average for the trait in control conditions ^15^:

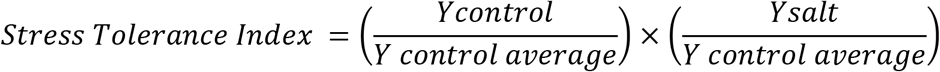

For each RIL, the control value was averaged across control replicates and an STI value was calculated for each salt treated plant. The STI value accounts for the overall performance of the population and compares each plants ability to maintain performance under stress conditions. Plants with large STI values indicate larger phenotypic values for a given trait and are often considered tolerant depending on the phenotype.

### Statistical analysis of phenotypic values

All statistical analyses were performed on RILs planted under control conditions, RILs planted under salt conditions, and on STI values. There were three biological replicates of each RIL under salt conditions, three biological replicates for STI values, and two biological replicates of each RIL under control conditions that were considered for QTL analysis. All statistics and graphing were completed using R version 4.0.2 ^44^.

Least square means for each phenotype in each condition (control, salt, STI) were calculated for every RIL. Normality was assessed using both a Shapiro-Wilk test in R and Q-Q plots from the *car* package version 3.0.10 ^45^. Traits that were not normally distributed were transformed (**Supplementary Table S1**). Transformed values were used in statistical tests and in QTL analysis. Correlations of phenotypes in each treatment (control or salt) were assessed via a Pearson’s Correlation analysis in R using the *PerformanceAnalytics* package 2.0.4 ^46^ (**Figure S1**).

To determine if there was a treatment effect, both a nonmetric multidimensional scaling (NMDS) analysis ^47^ and an analysis of variance (ANOVA) were performed. The NMDS ordination clustered individuals based on Bray–Curtis dissimilarity measures when all phenotypes were considered. The dimcheckMDS function in the *goeveg* package version 0.4.2 was used to generate stress values for each dimension; two dimensions were deemed appropriate. The NMDS was generated using the *vegan* package version 2.5.6 ^48^ in R. The NMDS was paired with an analysis of similarity (ANOSIM) to statistically test the ordination results. Specifically, the ANOSIM tested if plants were more similar within or between treatments (significance assessed at α = 0.05). An analysis of variance (ANOVA) was used to test if control and salt populations differed for individual phenotypes (significance assessed at α = 0.05).

### Genetic Map Construction and QTL analysis

The RIL population used in this study was generated as previously described in Govindarajulu et al. (2021). This population has been successful in identifying genes that are key regulators of tiller elongation in sorghum ^49^; however, in this study, advanced lines were included, therefore requiring a new genetic map. In summary, using high-quality nuclear DNA, the parent plants (*S. propinquum* and *S. bicolor*) were sequenced at 18x depth, while the RILs were sequenced at 2x depth. SNP data was aligned to the masked *Sorghum bicolor* reference genome version 3.1 ^50^. SNPs were analyzed with the GenosToABHPlugin in Tassel ver 5.0 ^51^ and loci were called as *S. propinquum* (A), *S. bicolor* (B), or *heterozygous* (H). The ABH formatted SNP data file was then used as input to SNPbinner ^52^, which calculated breakpoints ^49^. Breakpoints were merged if they were shorter than 0.2% of the chromosome length. After removing heterozygous bin markers and duplicate bin markers, the kosambi map function in R/qtl ^53,54^ was used to construct a high density genetic map.

QTL analysis was performed in R using the *qtl* package version 1.46.2 ^54^. QTL were first identified by a single interval mapping QTL model. Significant logarithm of the odds (LOD) peak scores were determined by comparing LOD peak scores after a 1,000 permutation test (α = 0.05) ^55^. If QTL were detected by interval mapping (IM), phenotypes were assessed via a multiple QTL model (MQM). The MQM tested for additional additive QTL, refined QTL positions, and tested for epistasis. Following MQM, a type III analysis of variance assessed the significance of fit for the final model, the proportion of variance explained, and the additive effect. QTL with a negative additive value indicated that the trait was negatively influenced by *S. bicolor* alleles, whereas a positive additive value indicated that the trait was positively influenced by *S. bicolor* alleles. Genes within a 1.0 logarithm of the odds (LOD) confidence interval for each QTL were identified (*Sorghum bicolor* ver. 3.1).

### Aquaporin Enrichment Analysis

We identified numerous genes that encode aquaporins (AQP) within 1.0 LOD of salt-specific QTL and hypothesized that there was an enrichment. To identify the number of AQP genes that would be expected to occur in an average sized QTL by random chance, 50 5 MB windows were randomly extracted from the *Sorghum bicolor* (version 3.1) genome in R. The starting position for each of the 50 samples was determined by first randomly selecting a chromosome and then randomly selecting a starting position on that chromosome. Chromosome selection was determined by multiplying the total number of chromosomes (2n = 10) by a randomly generated number [runif(1)] between 0 and 1, and truncating that number to an integer. This same process was repeated to determine the starting location on the chromosome; however, instead of multiplying a random number by the total number of chromosomes, the random number (between 0 and 1) was multiplied by the size of the corresponding chromosome. The result was again truncated and represented the starting location in the genome (in Mb). Genes within a 5 Mb window (5 Mb downstream from the starting location) were extracted (*Sorghum bicolor* version 3.1) for each of the 50 random samples. Because genes are unevenly distributed in the genome, the proportion of aquaporin genes per random sample or QTL was calculated (total number of AQP genes in region ÷ total number of genes in 5 Mb sample OR QTL). To statistically test if there was an enrichment of AQP genes within QTL, the proportion of AQP genes in the random samples were compared to the proportion of aquaporin genes in QTL with a two-tailed Student’s *t-*test using the t.test function in R. Significance was assessed at *P* ≤ 0.05.

## Results

### A high-density genetic map covers 10 *Sorghum* chromosomes with 1991 makers

The resequencing, SNP calling, and bin calling of the 177 RILs generated 4055 total bin markers (**Figure S2**). After removing duplicate markers, the map covered the 10 *Sorghum* chromosomes with 1991 markers (**Supplementary Table S2**) and was 913.71 cM in length (**Supplementary Table S3**).

### Overall plant health decreased in response to salt exposure

When comparing the RILs treated with salinity to control plants, we observed an overall decrease in plant health in response to salt exposure (**Figure S3**). Except for mortality, all phenotypes were significantly different between the control and salt treated populations (**Table 2**). Generally, the response to salt exposure was characterized as shorter plants with reduced live aboveground biomass, root biomass, total aboveground biomass, and total biomass (**Table 2, Supplementary Tables S4**). Further, plants had larger rank scores and more dead aboveground biomass (**Table 2, Supplementary Tables S4**). These significant differences between RILs planted under control conditions and salt conditions, in addition to the distinct and significant clustering of treatments in the NMDS analysis (p < 0.001, ANOSIM R=0.16)(**Figure S3**), are indicative of an overall decrease in plant health in response to salt.

**Table 2.**
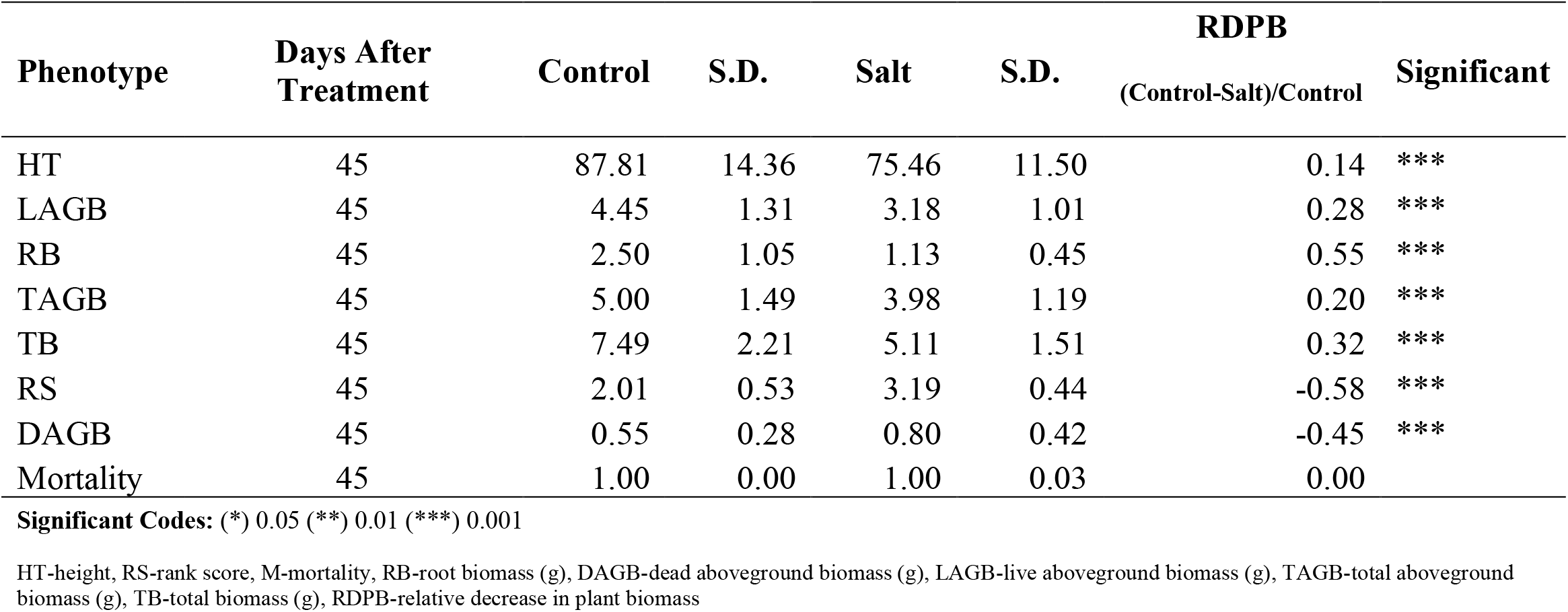
Summary of summary statistics and phenotypic averages for RILs in the control population and salt population. An analysis of variance (ANOVA) was used to test if there was significant variation in response to salt treatment. In response to salt exposure plants were shorter, had less live aboveground biomass, root biomass, total aboveground biomass, and total biomass. Additionally, in response to salt exposure plants had larger rank scores and more dead aboveground biomass. Mortality was not affected in response to salt treatment.

### In response to long-term salt exposure, QTL were identified as important regulators of salt tolerance

QTL analysis was performed on 1) RILs planted under control conditions, 2) RILs planted under salt conditions, and 3) on the stress tolerance index (STI) values ^15,56^. The individual QTL analyses performed in the presence and absence of salt were used to identify QTL that were shared. All genes within 1.0 LOD of QTL are listed in **Supplementary Table S5**.

#### Total biomass (TB)

In the absence of salinity, plants ranged from 1.98 g to 12.18 g of total biomass, with a mean of 7.49 g. In response to 75 mM NaCl, there was a decrease in the total plant biomass (0.66 g - 9.42 g with a mean of 5.11 g, **Table 2**, **Supplementary Table S4**). There were no QTL detected for total biomass when the control population was mapped. This is likely due to limited variation in architecture and biomass between *S. bicolor* and *S. propinquum* when plants are grown in space restricted pots (data not shown). When exposed to salinity, lines with *S. bicolor* alleles maintained significantly more biomass whereas lines with *S. propinquum* alleles accumulated 32% less biomass (**Table 2, Supplementary Table S4**).

Salt-specific QTL (qTB45_4.S and qTB45_4.STI) were detected on chromosome 4 (**Figure 1** and **Table 3**). The QTL had positive additive effects indicating that RILs with *S. bicolor* alleles in the 61.70 Mb - 68.41 Mb region on chromosome 4 have more overall biomass (live aboveground biomass, dead aboveground biomass, and root biomass) after long-term exposure to NaCl. The QTL detected when STI values were mapped (qTB45_4.STI) explained the greatest amount of phenotypic variation (PVE = 13.02) (**Table 3**). Of the genes within qTB45_4.STI, candidate genes were associated with aquaporins, stress response proteins, salt tolerant proteins, and transporters (**Supplementary Table S6**).

**Table 3.** QTLs identified in the RIL population using transformed least square means in control conditions, salt conditions, and with stress tolerance index values. The QTLs reported were identified when using Multiple QTL Mapping (MQM) in control conditions (0 mM NaCl), salt conditions (75 mM NaCl), and with stress tolerance index (STI) values. QTLs are named using the following system: q[Trait][DAT]_[Chr].[Treatment]

| Trait | Trt | QTL Name | DAT | Chr | Position (cM) | Bin (Max LOD) | Lod score | p-value | PVE | Additive | Start Mb | Peak Mb | End Mb |
| --- | --- | --- | --- | --- | --- | --- | --- | --- | --- | --- | --- | --- | --- |
| TB | S | qTB45_4.S | 45 | 4 | 63.7 | 62.29 | 3.73 | 3.90E-05 | 10.37 | 0.51 | 61.70 | 62.29 | 68.41 |
| TB | STI | qTB45_4.STI | 45 | 4 | 64.6 | 62.46 | 4.91 | 2.41E-06 | 13.40 | 0.09 | 62.06 | 62.46 | 64.38 |
| TAGB | S | qTAGB45_4.S | 45 | 4 | 83.4 | 67.29 | 3.40 | 8.65E-05 | 9.49 | 0.40 | 61.70 | 67.29 | 68.41 |
| TAGB | STI | qTAGB45_4.STI | 45 | 4 | 64 | 60.23 | 4.18 | 1.34E-05 | 11.55 | 0.09 | 61.91 | 60.23 | 67.44 |
| DAGB | C | qDAGB45_2.C | 45 | 2 | 72.7 | 65.42 | 4.91 | 1.63E-05 | 12.15 | -0.08 | 64.40 | 65.42 | 67.54 |
| DAGB | C | qDAGB45_9.C | 45 | 9 | 69.1 | 56.32 | 3.98 | 1.32E-04 | 9.71 | -0.06 | 55.47 | 56.32 | 57.91 |
| DAGB | STI | qDAGB45_2.STI | 45 | 2 | 73 | 62.28 | 4.21 | 1.24E-05 | 11.63 | -0.20 | 13.85 | 62.28 | 67.54 |
| LAGB | C | qLAGB45_4.C | 45 | 4 | 71.5 | 64.27 | 3.32 | 1.05E-04 | 9.22 | 0.45 | 62.17 | 64.27 | 67.29 |
| LAGB | STI | qLAGB45_4.STI | 45 | 4 | 73 | 63.41 | 4.22 | 1.22E-05 | 11.65 | 0.10 | 62.06 | 63.41 | 67.49 |
| RB | STI | qRB45_4.STI | 45 | 4 | 64.6 | 62.46 | 4.13 | 1.53E-05 | 11.40 | 0.08 | 62.17 | 62.46 | 62.54 |
| RS | STI | qRS45_4.STI | 45 | 4 | 69.1 | 63.67 | 3.89 | 2.71E-05 | 10.77 | -0.19 | 62.46 | 63.67 | 63.95 |
| HT | C | qHT45_7.C | 45 | 7 | 57.7 | 59.01 | 4.72 | 2.50E-05 | 11.18 | -5.00 | 58.66 | 59.01 | 60.24 |
| HT | C | qHT45_9.C | 45 | 9 | 63.4 | 55.07 | 5.32 | 6.57E-06 | 12.71 | -5.46 | 54.37 | 55.07 | 56.91 |
| HT | S | qHT45_1.S | 45 | 1 | 108 | 75.72 | 5.17 | 9.14E-06 | 12.57 | -4.44 | 76.22 | 75.72 | 80.57 |
| HT | S | qHT45_7.S | 45 | 7 | 63.5 | 60.17 | 4.95 | 1.51E-05 | 11.99 | -4.25 | 59.01 | 60.17 | 61.46 |
| HT | STI | qHT45_1.STI | 45 | 1 | 107.8 | 77.00 | 5.45 | 3.37E-03 | 10.63 | -0.04 | 75.67 | 77.00 | 80.57 |
| HT | STI | qHT45_4.STI | 45 | 4 | 82.2 | 66.96 | 4.66 | 1.16E-02 | 9.00 | 0.05 | 66.42 | 66.96 | 67.49 |
| HT | STI | qHT45_7.STI | 45 | 7 | 63.5 | 60.17 | 7.16 | 1.93E-04 | 14.34 | -0.05 | 59.01 | 60.17 | 60.24 |
| HT | STI | qHT45_9.STI | 45 | 9 | 63.4 | 55.07 | 4.43 | 1.66E-02 | 8.51 | -0.04 | 54.70 | 55.07 | 56.68 |

DAGB-dead aboveground biomass, LAGB-live aboveground biomass, RB-root biomass, RS-rank score, HT-height, TB-total biomass, TAGB-total aboveground biomass; Trt-treatment, STI-stress tolerance index, C-control, S-salt; P-*Sorghum propinquum*, B-*Sorghum bicolor*; DAT-days after treatment; Chr-chromosome, PVE-percent variance explained

**Figure 1.**
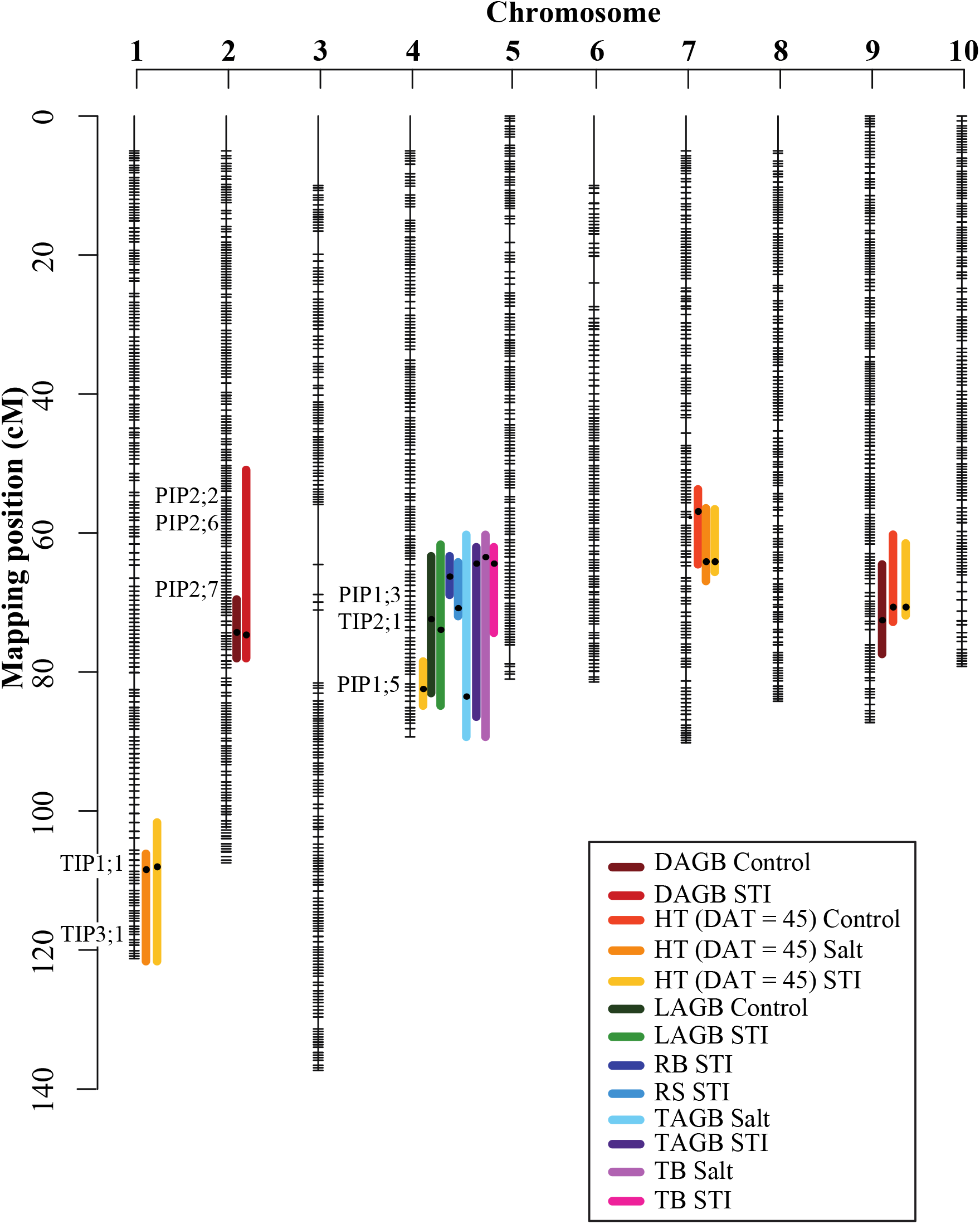
A genetic map with the QTL locations from 177_F3:F5_ *Sorghum RILs*. The empty spaces on each chromosome are regions that were removed because bins were either heterozygous or neighboring markers were identical (duplicate markers). The genetic map position is shown on the y-axis. Horizontal lines represent bins used as markers. Colored vertical lines show the position of each QTL for each trait in control conditions, salt conditions, or when STI values were mapped.

#### Total aboveground biomass (TAGB)

Plants grown in control conditions accumulated an average of 5.00 g of total aboveground biomass (1.34 g - 8.56 g), whereas in response to salt treatment there was a 20% decrease (average = 3.98 g) (**Table 2, Supplementary Table S4**). Similar QTL were detected when the total biomass and the total aboveground biomass were mapped, as would be expected given the high correlation between these two measures (**Supplementary Figure S1**). The same two QTL (qTB45_4.S and qTB45_4.STI) were detected in the salt population (qTAGB45_4.S) and when STI values (qTAGB45_4.STI) were mapped (**Figure 1**). Therefore, the same candidate genes identified in the total biomass QTL were also identified for total aboveground biomass.

#### Dead aboveground biomass (DAGB)

Dead aboveground biomass ranged from 0.06 g to 1.61 g (mean of 0.55 g) for plants grown under control conditions, whereas when plants were grown under salt conditions, there was 45% more dead aboveground biomass (**Table 2**, **Supplementary Table S4**). Salt sensitive genotypes, such as S. *propinquum* ^37^, accumulate more dead aboveground biomass compared to genotypes that are tolerant. When dead aboveground biomass was mapped, two QTL were detected on chromosome 2 (qDAGB45_2.C and qDAGB45_2.STI) and one QTL was detected on chromosome 9 (qDAGB45_9.C) (**Figure 1**). RILs with *S. propinquum* alleles positively influenced the amount of dead aboveground biomass. Of the genes within qDAGB45_2.STI, candidate genes were associated with aquaporins, sodium transporters, potassium transporters, salt tolerant proteins, and leaf senescence (**Supplementary Table S6**).

#### Live aboveground biomass (LAGB)

Live aboveground biomass for plants grown under control conditions ranged from 1.20 g to 6.95 g with an average of 4.45 g (**Supplementary Table S4**). In response to salt treatment, there was an average decrease of 28% in live aboveground biomass with a range from 0.32 g to 6.22 g and a mean of 3.18 g (**Table 2**). Two QTL were detected (qLAGB45_4.C and qLAGB45_4.STI) on chromosome 4 (**Figure 1**) and both QTL had positive additive effects, indicating that *S. bicolor* alleles positively influence live aboveground biomass. The QTL detected when STI values were mapped (qLAGB45_4.STI) explained 11.65 percent of phenotypic variation (**Table 3**).

#### Root biomass (RB)

Root biomass of plants grown in control conditions ranged from 0.48 g to 6.38 g with a population average of 2.50 g; however, plants grown in saline conditions accumulated an average of 55% less root biomass (**Table 2**, **Supplementary Table S4**). A single QTL was detected when STI values were mapped (qRB45_4.STI) (**Figure 1**). qRB45_4.STI explained 11.40 percent of the phenotypic variation and had an additive effect of 0.08, indicating that individuals with *S. bicolor* alleles in this region positively influenced root biomass.

#### Rank score (RS)

Rank score was a qualitative measure used to describe overall plant health (**Table 1**). There was a 58% average increase in rank score in response to treatment, indicating that there was a decrease in plant health (**Table 2**). Plants grown in control conditions had an average rank score of 2.01 (0.72−3.20) indicating that most of the individuals were beginning to show signs of leaf tip curling. In contrast, the average rank score of plants treated with salt was 3.19 with a range of 1.68 to 4.55 (**Supplementary Table S4**). This suggests that all individuals were displaying signs of stress, and some individuals stopped producing new leaves. RILs with a rank score near 1 or 2 in salt treated conditions are more tolerant of salinity. When STI values were mapped, a single QTL was detected on chromosome 4 (qRS45_4.STI). qRS45_4.STI is located at 62.46 Mb - 63.95 Mb with a peak near 63.67 Mb (**Figure 1**). Further, qRS45_4.STI explained 10.77 percent of the phenotypic variation and had a negative additive effect of 0.19, indicating that *S. bicolor* alleles are associated with overall better plant health in stress conditions. Of the genes located within qRS45_4.STI, there are several genes that encode aquaporins, ion channels, and chaperone proteins (**Supplementary Table S6**).

#### Height (HT)

The height of plants grown in control conditions ranged from 52.92 cm to 122.37 cm with a population average of 87.81 cm (**Supplementary Table S4**). In response to salt treatment, plants were an average of 16.38% shorter (**Table 2**, **Supplementary Table S4**). Eight QTL were detected for height (**Figure 1**). Three QTL (qHT45_7.C, qHT45_7.S, and qHT45_7.STI) were detected on chromosome 7 in approximately the same region (58.66 Mb - 61.46 Mb) and two QTL (qHT45_9.C and qHT45_9.STI) were detected on chromosome 9 in approximately the same region (54.37 Mb - 56.91 Mb) (**Table 3**). Since these five QTL were detected in plants grown in both control and treatment conditions, we suspect that genes in these QTL are related to plant architecture (e.g. height) rather than salt tolerance; however, two salt-specific QTL were detected on chromosome 1 (qHT45_1.S and qHT45_1.STI) and one on chromosome 4 (qHT45_4.STI)(**Figure 1**). Several candidate genes were identified, including genes associated with aquaporins, potassium transporters, and stress response proteins (**Supplementary Table S6**).

### Enrichment Analysis

A total of 4276 unique genes were located within 1.0 LOD confidence interval of the QTL identified in this study (**Table 3**, **Supplementary Table S5**). Of these, we observed 13 genes that encode aquaporins. To determine if this constitutes an enrichment of aquaporin genes, we used a two-tailed Student’s *t-*test to determine if the proportion of genes encoding aquaporins was higher in QTL compared to random samples of the genome. Of the 50 independent 5 Mb segments (34% of the genome), there were 8974 genes and six of the samples contained genes that encode aquaporins. As expected, results show that there was an enrichment of SbAQP genes in salt-specific QTL (mean = 0.0034) compared to 5 Mb random samples of the genome (mean = 0.0000, p≤.001).

## Discussion

In the present study, we screened 177 F_3:5_ RILs derived from a cross between the inbred *S. bicolor* (Tx7000; landrace durra) and its wild relative, *S. propinquum,* for performance-related traits in saline conditions. Because of sorghum’s importance in biofuel and forage production, salinity tolerance was assessed as the ability of the plant to maintain traits related to growth and performance in response to salt treatment. This tolerance can be achieved by various mechanisms including osmotic adjustment, Na^+^ exclusion from the aerial organs of the plant, and/or overall tissue tolerance; however, Na^+^ exclusion can also result from reduced Na^+^ uptake, increased Na^+^ extrusion to the roots and/or soil media, or increased retrieval from the shoot ^5,12–18^,^57^. Ultimately, each of these tolerance mechanisms results in the maintenance of plant vigor similar to plants grown in optimal conditions. In this study, we identified nineteen QTL from plants grown in control and salt conditions. Of the nineteen QTL, ten were either 1) detected only when STI values were mapped, 2) detected only from plants treated with 75 mM NaCl, and/or 3) explained more than ten percent of the phenotypic variation (**Table 3**), suggesting that these QTL are important in *Sorghum* salinity tolerance ^56^.

The data presented here, in combination with previous findings ^37^, collectively demonstrates that there is increased salt tolerance in *S. bicolor* compared to *S. propinquum*. For dead aboveground biomass, live aboveground biomass, total aboveground biomass, and total biomass, *S. bicolor* alleles were associated with tolerance. For example, the negative additive effect for the QTL detected for dead aboveground biomass indicates that *S. bicolor* alleles were associated with less accumulation of dead aboveground biomass. Similarly, the QTL detected for live aboveground biomass, total aboveground biomass, and total biomass all have positive additive effects indicating that *S. bicolor* alleles promote continued growth in stress conditions. It is important to note that, in optimal conditions, *S. propinquum* produces more aboveground biomass compared to *S. bicolor* ^49^. Therefore, in response to salt, the ability for lines with *S. bicolor* alleles to perform favorably supports the conclusion that *S. bicolor* possesses greater tolerance.

Because salinity stress is the product of both osmotic and ionic factors, their respective causes and consequences are often difficult to disentangle; however, these two stresses are often temporal in their action. When salts initially accumulate in the soil surrounding the root rhizosphere, the osmotic potential of the soil decreases, resulting in reduced water extraction by plant roots. Genotypes that are tolerant to osmotic stress exhibit the continuation of both above and belowground growth ^5^. Traits that we expected to be effected by osmotic stress include height, total biomass, and root biomass. Indeed, four salt-specific QTL (qHT45_4.STI, qTB45_4.STI, qTB45_4.S, qRB45_4.STI) were detected for these traits, co-localize on chromosome 4, and have positive additive effects indicating that *S. bicolor* alleles positively influence these traits. A comparison of our QTL with studies in maize reveal that two of the QTL (qHT45_1.S and qHT45_1.STI) identified in this study co-localize with salt-specific QTL in maize ^58^. Specifically, the QTL identified in maize were associated with greater shoot lengths, which is comparable to our height measurements in this study. Within these QTL, we identified genes that are associated with osmotic adjustment and water transport. For example, there were various genes whose products are involved in proline production, aquaporins, CDPKs (calcium-dependent protein kinases), sensing and signaling, cell division, Na^+^/Ca^2+^ exchanger, leaf senescence, early response to dehydration, heat shock proteins, vacuolar proton exchangers, potassium antiporters, and stress response proteins. These results suggest that genes associated with greater osmotic tolerance, as evidenced by maintained above ground biomass (height and biomass), are located within these QTL.

Following osmotic stress, the ion dependent phase occurs, and the most common phenotype associated with ionic stress is leaf necrosis. Therefore, we used dead aboveground biomass and rank score as a proxy for ionic toxicity. For qDAGB45_2.STI, *S. propinquum* alleles positively correlated with greater dead aboveground biomass, possibly because of increased Na^+^ accumulation in the aerial plant tissue. Similarly, qR.S45_4.STI also had a negative additive effect, indicating that *S. propinquum* alleles positively influenced the rank score, suggestive of greater susceptibility to ionic toxicity (**Table 1**). When comparing these QTL with QTL identified in other studies, we found that qDAGB45_2.STI co-localized with four QTL identified in maize ^58^. Within these QTL, we identified genes associated genes such as calcium-dependent protein kinases, LEA-like proteins, aquaporins, heat shock proteins, Na^+^/H^+^ antiporters, WRKY transcription factors, K^+^ uptake, and cation transporters (**Supplementary Table S5**).

Aquaporins are well known for their role in the transport of water and other neutral solutes ^59–65^. Emerging evidence indicates that some aquaporins are also capable of coupling water and ion transport, resulting in osmotic adjustment ^66^. Specifically, the *Arabidopsis* plasma membrane intrinsic proteins AtPIP2;1 (AT3G53420) and AtPIP2;2 (AT2G37170) have been shown to co-transport water and Na^+^, suggesting that they play dual roles in nutrient transport and osmotic adjustment ^66,67^. In the presence of salts, PIP2 is transported from the plasma membrane, resulting in significant reductions in root hydraulic conductance ^66,68,69^. In addition, the ionic conductance of AtPIP2;1 has been shown to be inhibited by divalent cations, specifically Ca^2+^, which is known to play an essential role in intracellular signaling in plants, particularly in response to abiotic stress ^62,66,70^. Therefore, AtPIP2;1 may constitute a mechanism of sensing and signaling during the salt stress response in plants. In *S. bicolor*, aquaporin transcript abundance has been shown to be affected by both ionic (salt) and osmotic (salt and drought) stress ^59^. Here, we identified 13 unique aquaporin genes in salt-specific QTL, which encode TIP1;1, TIP2;1, TIP3;1, PIP1;3, PIP1;4, PIP2;2, PIP2;6, PIP2;7, and PIP1;5 (**Supplementary Table S5**). Further, we identified the tandemly arranged Sobic.002G125000, Sobic.002G125200, Sobic.002G125300, and Sobic.002G125700, which encode four copies of SbAQP2;6 that lie within the salt-specific QTL (qDAGB_2.STI), and share ~85% sequence similarity with AtPIP2;1 (**Supplementary Table S5**). Additionally, seven genes identified in qDAGB45_2.STI encode plasma membrane intrinsic proteins (a family of aquaporins) in both sorghum and maize (**Supplementary Figure S6**). Given their presence in salt-specific QTL, demonstrated response to abiotic stress, and conservation among distantly related plant lineages (maize, sorghum, and *Arabidopisis*), aquaporins may play a critical role in maintaining water balance, controlling ion transport, and in sensing and signaling during the response to salinity stress in plants.

## Conclusions

In the present study, we detected numerous genes associated with sensing, signaling, and transport of Na^+^ in salt-specific QTL. Most interestingly, we identified numerous genes that encode aquaporins detected within salt responsive QTL. The results of this study provide insights into QTL important for each of the tolerance mechanisms (ionic, tissue, and osmotic tolerance). Additionally, this is the first study where individuals of each tolerance category (1-tolerant to ionic and osmotic stress; 2-tolerant to ionic stress but sensitive to osmotic stress; 3-sensitive to ionic stress and tolerant to osmotic stress; or 4-sensitive to ionic and osmotic stress) are identified in a common genetic background. Therefore, these findings and this population provide a foundation for future studies aimed at the dissection of the genetic basis of salinity tolerance.

## Data Availability Statement

Phenotype data and the binned genotype data used for QTL mapping can be found in Supplementary **Tables S7-S8**. All data necessary for confirming the conclusions of the article are present within the article, figures, and tables.

## Acknowledgements

We would like to acknowledge the WVU Genomics Core Facility, Morgantown WV for the support provided to help make this publication possible, and CTSI Grant #U54 GM104942 which in turn provides financial support to the Core Facility. The authors wish to thank Ryan Percifield for assistance during data collection, Dr. Stephen DiFazio and Dr. Sandra Simon for guidance in data analysis, Dr. Erin Sparks for reviewing this manuscript, and the West Virginia University Evansdale Greenhouse for supplying space. Additionally, we thank two anonymous reviewers and the associate editor for their insightful comments on this manuscript. This work was partially funded by the Eberly College of Arts and Sciences research award to Ashley N. Hostetler.

## Author Contributions

A.N.H and J.S.H designed experiments; A.N.H and J.S.H managed the project; A.N.H., prepared the samples; A.N.H. and R.G. lead the data analysis; A.N.H, and J.S.H wrote the manuscript with contributions from R.G.

## Supplemental Figures

**Figure S1. Pearson correlations on raw phenotypes and transformed phenotypes for control and salt populations at 45 days after treatment.** The distribution of each trait is shown on the diagonal. The bivariate scatterplot with the line of best fit for the corresponding traits is shown below the diagonal, and above each diagonal is the Pearson correlation coefficient and the associated p-value. Each significance level is indicated with an asterisk, where (***) corresponds to a p-value < 0.001, (**) corresponds to a p-value < 0.01, and (*) corresponds to a p-value < 0.05. Note: RB = root biomass; DAGB = dead aboveground biomass; LAGB = live aboveground biomass. (A) Control population 45 days after treatment (B) Transformed control data 45 days after treatment (C) Salt population 45 days after treatment.

**Figure S2. Sorghum genetic map after using a sliding window method to call bin markers as AA (***S. propinquum***), BB (***S. bicolor***), or AB.** (A) After the resequencing, SNP calling, and bin calling, 4055 total bins were detected across 10 sorghum chromosomes. (B) The 4055 total bins are illustrated on the x-axis and the 177 RILs are illustrated on the y-axis. The red regions illustrate bins that were called *S. propinquum* (AA); the green regions illustrate bins that were called *S. bicolor* (BB); the blue regions illustrate regions that were called heterozygous (AB).

**Figure S3. Non-metric multidimensional scaling (NMDS) analysis paired with an analysis of similarity (ANOSIM) reveals treatment clustering.** A NMDS paired with an ANOSIM reveals that individuals were more similar within a treatment than between treatments (ANOSIM R=0.16, p<0.001).

